# Context-invariant beliefs are supported by dynamic reconfiguration of single unit functional connectivity in prefrontal cortex

**DOI:** 10.1101/2023.07.30.551169

**Authors:** Jean-Paul Noel, Edoardo Balzani, Cristina Savin, Dora E. Angelaki

**Affiliations:** Center for Neural Science, New York University, New York City, NY, USA

## Abstract

Natural behaviors occur in closed action-perception loops and are supported by dynamic and flexible beliefs abstracted away from our immediate sensory milieu. How this real-world flexibility is instantiated in neural circuits remains unknown. Here we have macaques navigate in a virtual environment by primarily leveraging sensory (optic flow) signals, or by more heavily relying on acquired internal models. We record single-unit spiking activity simultaneously from the dorsomedial superior temporal area (MSTd), parietal area 7a, and the dorso-lateral prefrontal cortex (dlPFC). Results show that while animals were able to maintain adaptive task-relevant beliefs regardless of sensory context, the fine-grain statistical dependencies between neurons, particularly in 7a and dlPFC, dynamically remapped with the changing computational demands. In dlPFC, but not 7a, destroying these statistical dependencies abolished the area’s ability for cross-context decoding. Lastly, correlation analyses suggested that the more unit-to-unit couplings remapped in dlPFC, and the less they did so in MSTd, the less were population codes and behavior impacted by the loss of sensory evidence. We conclude that dynamic functional connectivity between prefrontal cortex neurons maintains a stable population code and context-invariant beliefs during naturalistic behavior with closed action-perception loops.

## Introduction

Adaptive behaviors require flexible computations abstracting away task features and goals from our immediate sensory context. This ability allows identical sensory input to yield different motor outputs under distinct contexts (i.e., context-dependence). Or conversely, for a single task to be accomplished under vastly different sensory conditions (i.e., context-invariance). We may, for instance, navigate to our colleague’s office in broad daylight by leveraging visual observations (e.g., optic flow), or toward our fridge in complete darkness by relying on acquired internal models – e.g., a cognitive map^1^ of our home, the size of our stride. The mechanisms subserving these flexible computations are not fully understood.

Two broad frameworks for context dependent computation have been proposed. First, brain-wide neuroimaging studies in humans^2-4^ have argued for “flexible hubs”. This work has highlighted the malleable nature of functional connectivity patterns, and the fact that these patterns change with task demands. This framework has primarily emphasized the role of the fronto-parietal network, with functional connectivity within this network being (i) the most flexible, and (ii) sufficient to identity which task is being undertaken^5^. The second approach has argued for a dynamical systems understanding. Namely, neurophysiological recordings in macaques^6-9^ coupled with population-level analyses have suggested that task context may either mold neural dynamics themselves^6^, or the initial condition of neural trajectories within a fixed dynamical system^7^. Of course, these frameworks are likely not entirely independent, with theory^10-12^ and nascent empirical observations^13-15^ suggesting that the statistical dependencies between neural responses (i.e., functional connectivity) facilitate context-invariant decoding, as well as restrict spiking activity to a low-dimensional^16^ and differentiable^17^ space.

The study of neural co-fluctuations and their relation to population dynamics and flexible behavior has, however, focused primarily on the representation of simple and static stimuli^18^, and on stereotyped behaviors^13-15, 18, 19^. Moreover, it has mostly examined a single neural node at a time^18-20^, and commonly relied on measurement techniques (e.g., fMRI, calcium imaging) that allow for sampling large neural populations at the expense of temporal resolution^13-15, 21^. Further, the relation between single cell functional connectivity and population dynamics supporting flexible behavior has received more theoretical than empirical evidence (see^6, 7, 9, 22-24^ for work employing artificial RNNs as “proof of principle” but less biological evidence). This is likely due to the difficulty in statistically quantifying unit-to-unit couplings in physiological data. Most vexingly, we do not understand how inter-neuronal correlations shape the dynamics supporting *naturalistic* behaviors; those unfolding over a protracted time period, within closed action-perception loops, and reliant not only on static percepts, but on dynamic *beliefs* regarding latent variables.

To close these gaps in knowledge, here we record singe cells in sensory, parietal, and frontal cortices while leveraging two key innovations. First, we developed a closed-loop task wherein macaques navigate in virtual reality and stop at the location of briefly visible targets^25^ – much akin to “catching fireflies.” Successful completion of this task requires animals to dynamically update their beliefs over a latent variable: the evolving distance to the (invisible) target. Importantly, this naturalistic task allows for rapid generalization^26^, and thus while animals were trained to path integrate by primarily accumulating optic flow signals (“high-density condition”, **Fig. 1A**), they may also be tasked with navigating in a visually impoverished environment (“low-density condition”, 50-fold decrease in optic flow density, see^27, 28^). Normatively, in this latter context, animals had to substantially rely on their internal models mapping joystick position to velocity in virtual space. Thus, animals performed a single task under two distinct computational contexts, defined by the relative weighting between sensory information and an acquired internal model (see^27 28^ for evidence in untrained human observers that a reduction in density in optic flow results in stronger reliance on a well-established slow speed prior).

**Figure 1.**
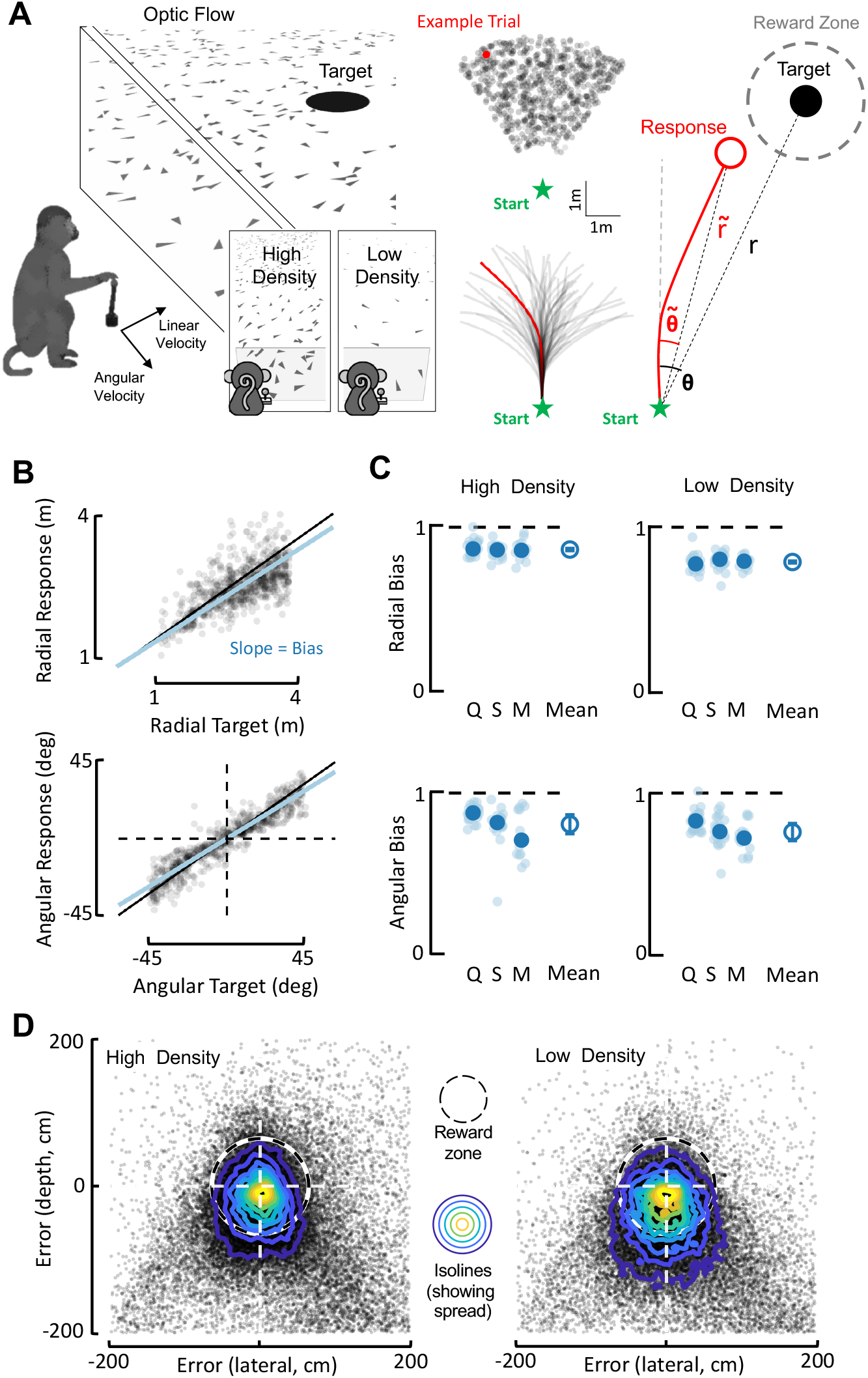
Macaques navigate in virtual reality to latent targets using path integration and an acquired internal model mapping joystick position to self-motion velocity. **A. Setup**. Monkeys use a joystick controlling their linear (max = 200cm/s) and angular (max = 90deg/s) velocity to navigate to the location of a briefly (300ms) presented target. Monkeys were trained with a high-density of ground-plane elements, thus having access to a self-velocity signal (motion across their retina) they integrated into an evolving estimate of position. During recordings, we additionally used a low-density condition, wherein monkeys had to primarily rely on an internal model mapping joystick position to virtual velocity. Targets were presented within a radial (1-4m) and angular (-45 to 45 deg relative to straight ahead) range, and we quantified performance by regressing radial (*r*) and angular eccentricity (*θ*) of responses and targets. **B. Example session**. Monkeys tended to undershoot targets in radial distance (top) and eccentricity (bottom). For each session, we fit a line and thus a slope of ‘1’ indicates no bias. Slopes smaller than 1 indicate undershooting. **C. Radial and angular biases during high- and low-density conditions**. Q, S, and M refer to three individual monkeys. Transparent small circles are individual sessions, and the larger opaque blue circle is the mean for each animal. The rightmost circle within each panel is the mean across animals, with error bars indicating ± 1 S.E.M. **D. Endpoint error and variance**. Scatter plot of endpoint errors along the depth (y-axis) and lateral (x-axis) dimension as a function of density. For visualization, data is pooled across all sessions and monkeys (high-density n = 44476 trials; low-density n = 43286 trials). Isolines showing 2 dimensional histograms demonstrate the increased spread (variance) of endpoints relative to target in the low-density condition.

The second innovation is technical. In natural behaviors no two trials are alike, and thus we cannot compute traditional noise correlations^29^ by averaging and extracting residuals. To address this challenge, we developed a Poisson Generalized Additive Model (P-GAM)^30^ to account for stimuli-driven responses (i.e., tuning functions) and estimate time-dependent and signal-independent coupling filters between neurons (i.e., the likelihood of a neuron firing given the firing of other neurons). Critically, our estimates include a confidence intervals for all parameters and thus allow for individually determining their significance in explaining the data (see^31, 32^ for a similar non-Bayesian approach without the ability for inferential statistics). The overall (i.e., integrated over time) strength in coupling between two units is akin to a traditional noise correlation metric, in that signal-independent co-fluctuations are indexed. Further, establishing time-resolved coupling filters allows us to examine the finer grain dynamics in statistical co-fluctuations between neurons. This technique recapitulates known properties of noise correlations, such as its dependency on the distance between neurons ^33^ and tuning similarity ^20, 34^ (see **Fig. S1**, also see^35^ for a detailed description of spiking responses and spike-LFP phase coupling during optic-flow guided navigation, i.e., the “high-density” condition). Furthermore, establishing time-resolved coupling filters allows us to examine the finer grain dynamics in statistical co-fluctuations between neurons, unavailable with other methods.

Our results demonstrate a decrease in the relative weighting of sensory evidence vis-à-vis an acquired internal model during navigation, accompanied by a strengthening of fixed coupling filters in sensory cortex (i.e., a gain modulation) and a dynamic reconfiguration of the finer-grain, time-resolved coupling in parietal and frontal cortices (i.e., remapping). In contrast, population dynamics were context-invariant in parietal and prefrontal cortices, but not in sensory cortex. Critically, the more the fine-grain statistical co-fluctuations reconfigured across contexts in prefrontal cortex, but not in parietal cortex, the more stable were its population dynamics and the less was the animals’ performance impacted by the loss of sensory evidence. Highlighting the central role of noise correlations in engendering a stable population code in prefrontal cortex, destroying the statistical dependencies between neurons in this area, but not in parietal or sensory cortices, eliminated this population’s ability for cross-context decoding of the key latent variable, distance to target.

## Results

### A single behavior under different computational demands

Prior work has studied the single neuron correlates of flexible behavior by having macaques alternate between two different tasks (e.g., color or motion discrimination^6, 36^). Here, instead, we employ a single task – navigate to target – that may be accomplished under different computational constraints. We have macaques (Q, S, and M) use a joystick (linear and angular velocity control) to path integrate in virtual reality and stop at the location of a briefly presented (300ms) target (**Fig. 1A**). Across trials, targets were randomly distributed within a radial (1m to 4m) and angular (-45deg to 45deg) range (uniform distributions, independently sampled). The virtual environment was solely composed of ground-plane elements, which flickered (250ms lifetime) and thus provided an optic flow signal but no landmark information. On a trial-by-trial basis we manipulated the density of these ground-plane elements, which were either in a “high-density” (5.0 elements/m^2^) or “low-density” (0.1 elements/m^2^) configuration (50-fold change).

Task performance was quantified by expressing the monkeys’ trajectory endpoints and target locations in polar coordinates, with an eccentricity from straight-ahead (*θ*) and a radial distance (*r*, **Fig. 1A**, rightmost). **Figure 1B** shows radial (top; *r* vs. 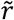 slope = 0.90; R^2^ = 0.55) and angular (bottom; *θ* vs. 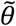 slope = 0.95; R^2^ = 0.78) responses as a function of target location for an example session (see *Methods* for detail). Most importantly, across all animals (n = 3) and sessions (n = 82), macaques were accurate in navigating to target, both during high-(regression slope, mean ± S.E.M., radial = 0.85 ± 0.04; angular = 0.82 ± 0.14) and low-density conditions (radial = 0.79 ± 0.04; angular = 0.77 ± 0.12; contrast of high vs. low density separately for each animal and displacement dimension, all p > 0.13, **Fig. 1C**). Thus, macaques were able to adaptively estimate their evolving distance to target regardless of the density of optic flow elements and the required computation.

While the location of mean endpoints did not change across densities (**Fig. 1B** and **C**), the variance of these endpoints did (distance to target S.E.M. computed on each session, high density, 0.19 ± 0.08; low density, 0.25 ± 0.08; p = 0.044, **Fig. 1D**). This resulted in a mean decrease of 4.41% of rewarded trials in the low-density condition relative to the high-density condition (fraction trials rewarded, high density, 0.60 ± 0.01; low density, 0.55 ± 0.01). More importantly, under a Bayesian framework, a change in the variance of trajectory endpoints is precisely what is expected when sensory likelihoods widen (i.e., decrease in sensory reliability) and priors are unbiased – as when animals were trained on the task. Instead, when observers are not trained on the task (feasible in humans but not in macaques), they have a biased slow-speed prior^27, 37^ and thus show both an increased variance^28^ and overshooting^27^ of targets during low-density optic flow path integration. Together, these results demonstrate that macaques trained to navigate by integrating optic-flow velocity cues were able to generalize this behavior and navigate by more heavily relying on an acquired internal model when sensory evidence was reduced.

### Context differentially impacts neural statistical structure within sensory, parietal, and prefrontal cortices

We recorded single unit spiking activity simultaneously in sensory (i.e., dorsomedial superior temporal area, MSTd, 240 neurons), parietal (area 7a, 2647 neurons) and prefrontal cortices (dorsolateral prefrontal cortex, dlPFC, 445 neurons) to probe how cortical nodes, and in particular their statistical dependencies, support a population code and adaptive beliefs despite changing environments. **Figure S2** shows peri-event time histogram (PETH) for example neurons in each brain area, as a response to different events (e.g., target or movement onset), and across density conditions. To quantitatively summarize these responses, we fit a P-GAM^30^ to account for stimulus-driven responses, as well as for elements of global neural dynamics (i.e., the ongoing phase of LFP within theta, alpha and beta ranges). Importantly, the encoding models also included hundreds of potential coupling filters capturing unit-to-unit time-resolved statistical regularities (exact number being session specific**; Fig. 2A**, highlighted in orange). The P-GAM accounted well for spiking activity (average pseudo-r^2^ = 0.072, ∼1.5 to 2 times better than traditional Generalized Linear Models^38^, see **Fig. S3A** for example tuning functions, both as raw binned responses and as estimated by the P-GAM. Also see^35^ for a detailed characterization of signal correlation, including mixed selectivity^39, 40^, during the high-density condition).

**Figure 2.**
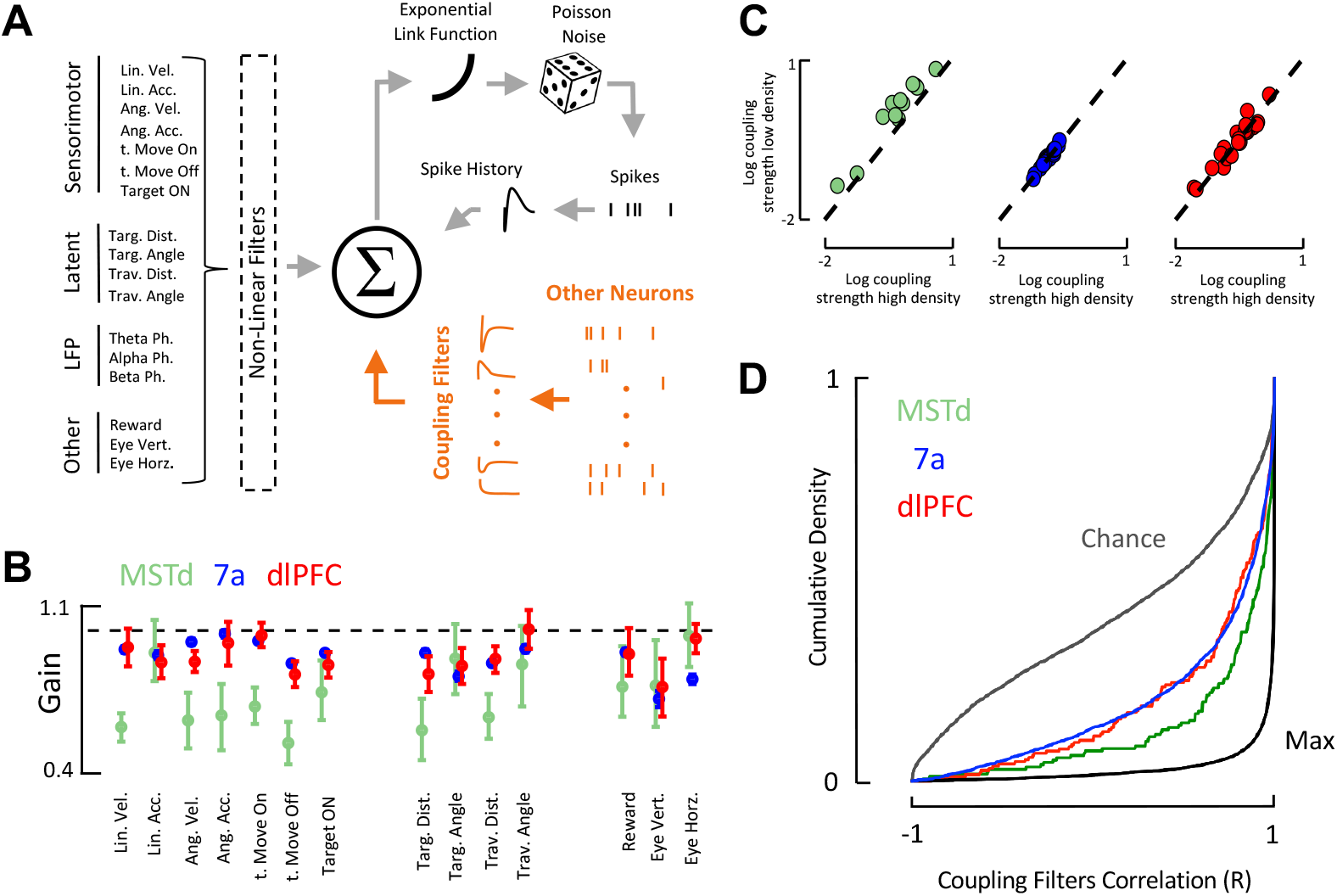
Estimation of coupling filters and their remapping with optic flow density. **A. P-GAM model**. The encoding model included 17 analog or digital task-variables, as well as spike history and coupling filters (in orange for emphasis). **B. Tuning function gain**. Gain in tuning functions in the low-density condition relative to the high-density optic flow condition. A value under ‘1’ (dashed black line) shows that the amplitude of the tuning function was reduced in the low-density condition relative to the high-density. Error bars are ± 95%CI. **C. Coupling filter strength**. For all pairs of neurons that were coupled both in the high- and low-density conditions, we quantified the strength of their coupling (i.e., norm of betas) and averaged within sessions. Coupling strength increased in the low-density condition in MSTd (top; green) but was unchanged in area 7a (middle; blue) and dlPFC (bottom, red). Each dot is a session. **D. Coupling filter shape stability**. Cumulative density of the coupling filters correlation coefficient (r^2^) between low- and high-density conditions for all three brain regions. Chance and ceiling-levels were determined by permutation. Coupling filters significantly changed across optic-flow density condition in dlPFC and 7a.

Regarding signal correlations, the fraction of neurons tuned to different task variables was not different across high- and low-density conditions (all p > 0.25, **Fig. S3B**). To quantify the impact of the different optic flow densities on neural tuning gain, we correlated the tuning functions in each of these conditions (**Fig. S3C**), given they were both significant (87.8% of cases, Kruskal-Wallis contrasting r-values across areas, p = 0. 24). The slope of this correlation quantifies a change in the gain of neural responses driven by task-relevant stimuli. While we observed significant gain modulation in all brain regions (all p < 0.005), the effect size was considerable in MSTd (Cohen’s d = 0.21) and negligible in the other areas (Cohen’s d < 0.06 in 7a and dlPFC; **Fig. 2B**, also see examples in **Fig. S2**). Neural responses were less prominent in MSTd (i.e., gains < 1) during the low than the high-density condition, suggesting that while signal correlations were driven by environmental input in MSTd, they were less so in 7a and dlPFC.

Regarding noise correlations, the fraction of units functionally coupled, either within (all p>0.88) or across areas (all p>0.76) was unchanged across the different environments (**Fig. S3D**, 94.1% of couples retained their coupling status across density conditions). On average, 20.4% of within-area coupling filters were significant, while only 1.08% of across area coupling filters were. Thus, for statistical power, all subsequent analyses focused on the within-area coupling filters.

The coupling strength (i.e., area under the coupling filter, collapsed across time) did not change with optic flow density within 7a or dlPFC (both p > 0.67, both Cohen’s d < 0.05). On the other hand, coupling strength within MSTd increased during the low-density manipulation (p<0.005, paired t-test, Cohen’s d= 0.51; **Fig. 2C)**, as if attempting to offset the impact of the unavailability of sensory evidence and the consequent reduced gain in neural tuning in MSTd (**Fig. 2B**). The correlation between change in tuning gain and coupling strength across sessions in MSTd showed a weak trend but was not significant (r = -0.23, p = 0.19).

To examine the finer grain temporal structure of neural co-fluctuations, we correlated coupling filters across density conditions (r-values being independent of gain, see **Fig. S4** for example coupling filters in each area and across density conditions). We also correlated coupling filters estimated across different recording sessions (i.e., not simultaneously recorded) and as estimated on odd vs. even trials (within a single density), to respectively approximate chance and maximum correlation levels. This analysis showed that the temporal structure of stimuli-independent neural co-fluctuations partially remapped with context in all areas (greater than chance, smaller than max), but did so less in MSTd than in 7a and dlPFC (Kruskal-Wallis contrasting areas, p = 0.002; post-hoc Holm-Sidak corrected contrasts, MSTd vs. 7a, p = 0.003; MSTd vs. dlPFC, p = 0.02; 7a vs. dlPFC, p = 0.76, **Fig. 2D**). Together, these results suggest that while behaviorally the animals were able to maintain adaptive beliefs over their distance to target regardless of sensory environment (**Fig. 1**), the fine grained statistical dependencies between neurons, particularly in area 7a and dlPFC, dynamically remapped with the changing computational demands (**Fig. 2**). Next, we questioned how a dynamically changing neural code may engender a context-invariant belief supporting adaptive behavior.

### Stability of population code and relation to dynamic coupling filter remapping

We conjectured that the dynamic reconfiguration of fine-grained statistical dependencies between neurons may support flexible behavior by rendering population dynamics invariant to the changing sensory input. And vice-versa, without coupling filters remapping, distinct population dynamics would emerge under different contexts (due to the changing sensory input). Accordingly, we hypothesized that 7a and dlPFC, but less so MSTd, would demonstrate context invariant population codes – despite (or even due to) their changing unit-to-unit couplings.

In a first step, we examined global population dynamics by principal component analysis (PCA). We binned trials according to target location (**Fig. 3A**, top row, overhead view) and optic-flow density (HD = “high-density”, LD = “low density”). We then time-warped, averaged across trials, and visualized the first two PCs. The results show that while population dynamics changed with density in MSTd, they did not in 7a and dlPFC (**Fig. 3A**, second through fourth row. See **Fig. S5** for quantification beyond the first 2 PCs and result showing that in higher-dimensions the population code in dlPFC, but not 7a, does change with optic-flow density).

**Figure 3.**
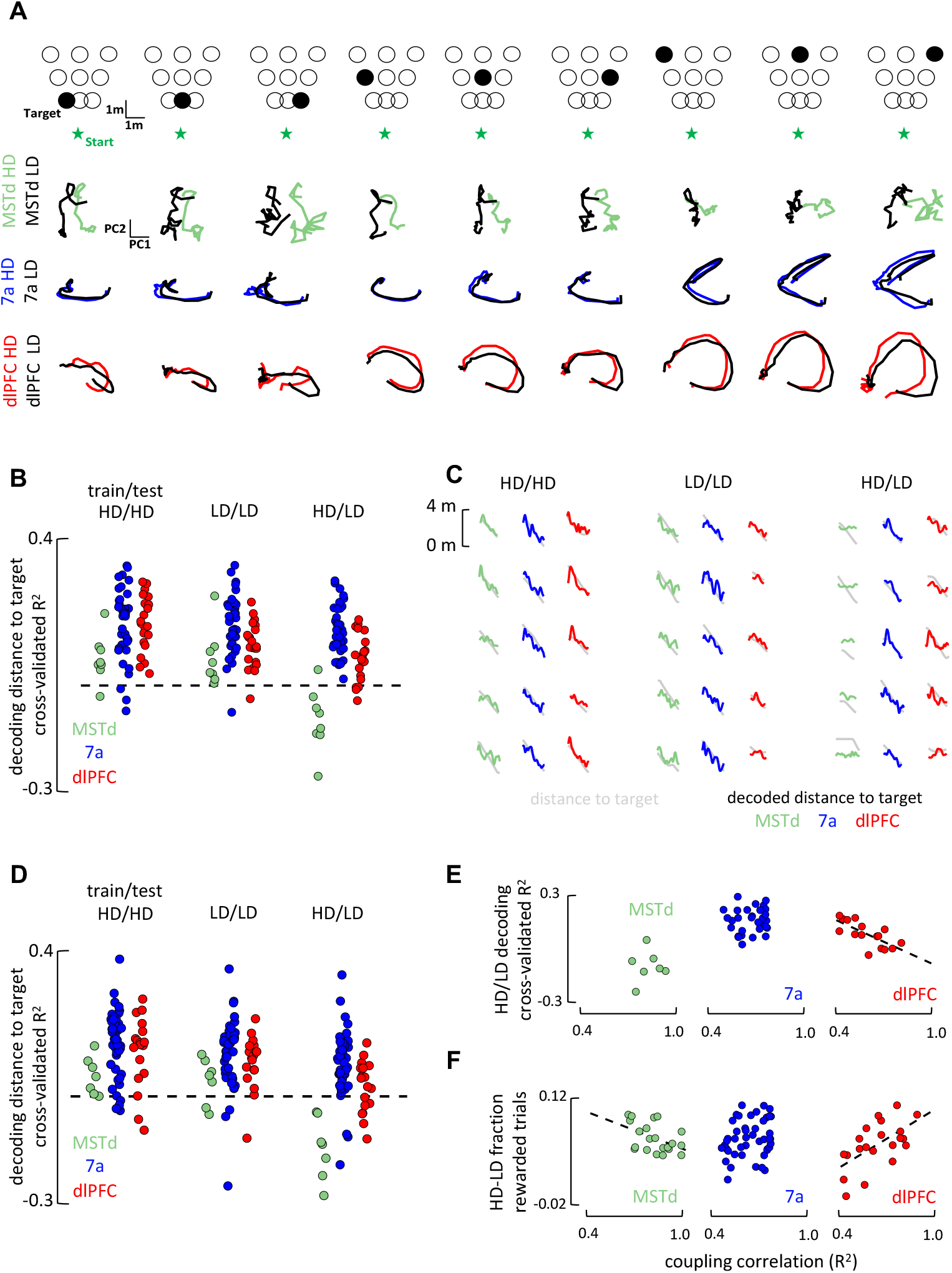
Population codes and their relation to coupling filter remapping and behavior. A. Illustration of task-unaligned latent neural dynamics. Trials were categorized according to their target location (top: 9 bins, from near-to-far and from leftward-to-rightward) and density condition (bird’s eye view, first row). Second, third, and fourth row respectively show the averaged latent trajectory in 2D for MSTd (green), 7a (blue), and dlPFC (red), both during high (colored) and low (black) density conditions, and according to target location (first row, 9 columns). **B. Decoding of the distance to target based on neural population dynamics**. For sessions with at least 15 neurons per brain area, we decode the distance to target while training and testing either within condition (High-density/High-density, Low-density/Low-density) or across (High-density/Low-density). Across condition decoding was possible in 7a (blue) and dlPFC (red), but not MSTd (green). Dots are individual sessions, dashed black line is cross-validated R^2^ = 0. **C. Five example trials decoding distance to target**. Observed distance to target (gray) is overlayed with the decoded distance in MST (green), 7a (blue), and dlPFC (red) when trained and tested on the same condition (HD/HD, LD/LD), or when the decoder was asked to generalize (HD/LD). **D. Decoding of the distance to target based on populations where signal correlations were left intact but noise correlations were destroyed**. Follows the formatting from **B** and shows the inability for dlPFC to generalize decoding of distance to target when noise correlations are eliminated. **E. Relation between stability of the fine-grain neural co-fluctuations and ability of the decoder to generalize**. For sessions with at least 15 neurons per area and at least 5 units significantly coupled with at least one other unit, we performed a correlation between their coupling filters stability (x-axis, R^2^) and decoder performance in HD/LD. **F. Relation between stability of the fine-grain neural co-fluctuations and session-by-session changes in behavioral performance with optic flow density**. For sessions with at least 5 units significantly coupled with at least one other unit, we correlated the difference in fraction of rewarded trials (HD-LD) and correlation between coupling filters across conditions. The more coupling filters remapped in dlPFC (lower R^2^), the less was performance impacted by the loss of sensory evidence in the low-density optic flow condition. Instead, the more stable where the coupling filters in MSTd (higher R^2^) the less was performance impacted by the loss of sensory evidence.

Next, to quantitively test the hypothesis that the manifold coding for distance to target – as opposed to population dynamics as a whole (**Fig. 3A**) – would remain stable in 7a and dlPFC, we used a linear population decoder (Lasso regression, 5-fold cross-validation) to estimate the animal’s evolving distance to target. Most importantly, to test for the stability of this task-relevant manifold, we assessed the ability of these decoders to generalize across density conditions (i.e., train on high-density and test on low-density). Results showed that while all areas were able to decode distance to target when trained and tested within a single context (**Fig. 3B**, left and center, cross-validated R^2^>0, all p < 0.029), the decoders generalized across contexts in area 7a and dlPFC (CV R^2^>0, respectively, p = 4.19 x10^-15^, and p = 2.57 x10^-5^), but not in MSTd (CV R^2^<0, p = 0.01, **Fig. 3B**, rightmost, see **Fig. 3C** for example single trial decoding of distance to target). This suggests the existence of a similar population readout for distance to target across contexts in 7a and dlPFC, but not MSTd.

Three pieces of evidence suggest that the stability of the manifold coding for distance to target in dlPFC, but not that in 7a, is related to the dynamic remapping of temporally-resolved unit-to-unit couplings across densities. First, artificially eliminating the statistical dependencies between neurons (see *Methods*) abolished the ability of dlPFC (CV R^2^>0, p = 0.84), but not area 7a (CV R^2^>0, p = 1.17 x10^-8^), in decoding distance to target across densities (HD/LD decoding, **Fig. 3D**). Eliminating these neural co-fluctuations did not change global neural dynamics as quantified by the first two PCs, which were primarily driven by signal and not noise correlations. Second, there were strong and specific session-to-session correlations between the stability of coupling filters in dlPFC and the ability to decode across contexts. That is, the more unit-to-unit couplings remapped within dlPFC in a session, the better was this session’s ability to decode distance to target across contexts (r = -0.71, p = 0.0018; **Fig. 3E**). This was not true for area 7a, where coupling filter stability did not correlate with decoding generalizability (r = 0.08, p = 0.65). This was also not true for signal correlations, with the stability of tuning functions across densities not correlating with decoding generalizability in 7a (p = 0.21) or dlPFC (p = 0.12). Third, the degree of remapping of neural co-fluctuations in dlPFC (and its stability in MSTd) were related to behavioral performance. Indeed, we observed that the more coupling filters remapped in dlPFC (r = 0.59, p = 0.004), the less was behavior impacted by the loss of sensory evidence in the low density condition (**Fig. 3F**). Interestingly, we noted the opposite effect in MSTd, with more stable coupling filters resulting in behavioral performance being less impacted by the loss of sensory evidence (r = -0.47, p = 0.030). There was no relation between behavioral performance and coupling filter stability in area 7a (r = 0.18, p = 0.22). Lastly, demonstrating the specificity of these effects, there was no relation between decoder generalizability or behavioral performance and changes in coupling strength (all p > 0.58).

Together, these results suggest a context-invariant population code – both global and aligned to the critical task variable, distance to target – in area 7a and dlPFC. These areas are also those demonstrating significant remapping of unit-to-unit couplings with context (**Fig. 2D, Fig S4**). Interestingly, the results suggest a relation between the stability of the task-relevant population code and coupling filters remapping in dlPFC but not in 7a.

## Discussion

We leveraged a task allowing for rapid generalization^26^ and a naturalistic interplay between sensory input, internal neural dynamics, and motor output in an attempt to bridge the gap between functional connectivity, population codes, and flexible behavior. To do so, we also leveraged a statistical methods innovation^30^ to estimate time-resolved coupling filters and their statistical contribution to a neuron’s spiking activity. This approach opens a window of opportunity above and beyond the traditional estimates of noise correlation^29, 34^, which are not time-resolved and cannot be easily applied to complex naturalistic tasks.

The most remarkable observation is that co-fluctuation of neural activity changed differently within sensory and parietal/frontal cortices when animals were tasked to further rely on acquired internal models, as opposed to sensory evidence. While MSTd seemingly attempts to rigidly extract environmental signals (i.e., increased strength in noise correlations albeit with fixed dynamics), the fine-grain statistical dependencies between neurons in 7a and dlPFC flexibly reconfigured under different contexts. This observation is reminiscent of a wealth of human neuroimaging studies^2-5^ emphasizing the flexible remapping of the fronto-parietal network with changing task demands. Our results also concord with previous recordings in macaques showing context or task dependent modulations in spiking activity in prefrontal^6-8^ and parietal^42^ cortices, but not extrastriate^42^ cortex. However, at difference from those studies, we must note that here we do not observe context-dependent differences in average firing rates (i.e., tuning functions) in 7a and dlPFC, but in the pairwise interactions of neurons within these areas. To the best of our knowledge, no prior primate studies have assessed if and how time-resolved coupling filters change with context across cortical areas.

The relation between remapping coupling filters, population codes, and flexible behavior also differed between dlPFC and 7a. Seemingly, dlPFC flexibly reconfigured to maintain an adaptive latent representation (i.e., “computation through dynamics^43^”). This is supported by the fact that the degree to which dlPFC remapped across contexts in different sessions related to the stability of the distance to target read-out manifold, as well as to behavioral performance. Further, eliminating the statistical dependencies between neural responses in dlPFC abolished the ability of this area for cross-context decoding. Area 7a similarly exhibited a context-invariant population code, yet this ability appeared unrelated to the remapping of coupling filters. This latter observation suggests that the context-invariant population dynamics in area 7a are potentially downstream (i.e., inherited) from those in dlPFC, that this area is less driven by visual flow (or the lack thereof) than it is by body-related (e.g., vestibular, proprioceptive, motor) or decisional signals (see^35, 44, 45^ for evidence to this regard), and/or that relative to dlPFC, population responses in 7a were less driven by noise correlations and more by signal correlations and/or LFP. Fittingly, we observe that higher PCA dimensions (i.e., accounting for less variance and thus less driven by task inputs and more by noise correlations) varied with context in dlPFC but not in 7a (**Fig. S5B**). Similarly, in prior work^35^ we have demonstrated that area 7a is strongly driven by sensorimotor variables and the ongoing phase of LFP in different frequency bands. Thus, while the population code in area 7a was context-invariant, this appears to be less dependent on the structure of unit-to-unit statistical co-fluctuations.

Mechanistically, some^6^ have argued that a change in context results in an alteration in recurrent dynamics. Others^7^ have assumed that connectivity does not change at this time scale, and thus neither can the underlying neural dynamics. Instead, according to this latter view, changes in context ought to alter the initial condition or input into the dynamical system. At first glance, it would appear that our results support the former interpretation, given that we report changes in functional connectivity. However, we must highlight that given the closed-loop nature of our task, input to the system are state-dependent and continuously changing. This renders challenging, if not impossible, the teasing apart of the relative contributions of internally generated dynamics from external inputs^46^. More importantly, we consider this a feature rather than a bug. In real-world behaviors, our brains, bodies, and environments ought to be considered a single dynamical system, so that teasing apart the role of internal neural dynamics and inputs from the environment may be a false dichotomy. Instead, we highlight that when faced with different environments posing different computational challenges, the brain appears to adopt a two-pronged approach. Sensory areas may not change their apparent functional connectivity, while frontal areas dynamically remap. We propose that this approach may allow for a compromise between faithfully reflecting our sensory environment (i.e., representation in sensory areas) and not completely changing our interpretation of the world around us (i.e., coding of latent variables or beliefs in prefrontal cortices) given stochastic fluctuations in environmental input.

## Materials and Methods

### Animals and Animal Preparation

Three adult male rhesus macaques (Macaca Mulatta; 9.8-13.1kg) were studied. We analyzed behavioral and neural data from 27 sessions in monkey Q, 38 sessions in monkey S, and 17 sessions in monkey M. These same sessions have been reported on before^35^, but only the “high-density” trials. The “low-density” data and their comparison to the “high-density” condition is novel. All animals were chronically implanted with a lightweight polyacetal ring for head restraint. Further, for acute recordings, animals were outfitted with a removable grid to guide electrode penetrations. Monkeys were trained via standard operant conditioning to path integrate to the location of briefly visible targets (“fireflies”) by accumulating optic flow velocity signals caused by active self-motion and thus motion of optic flow elements across their retina. All surgeries and procedures were approved by the Institutional Animal Care and Use Committee at Baylor College of Medicine and New York University, and were in accordance with National Institutes of Health guidelines.

### Experimental Setup

Monkeys were head-fixed and secured in a primate chair. A 3-chip DLP projector (Christie Digital Mirage 2000, Cypress, CA, USA) rear-projected images onto a 60 x 60 cm screen that was attached to the front of the field coil frame, 32.5 cm in front of the monkey. The projector rendered stereoscopic images generated by an OpenGL accelerator board (Nvidia Quadro FX 3000G). To navigate, the animals used an analog joystick (M20U9T-N82, CTI electronics) with two degrees of freedom to control their linear (max = 200cm/s) and angular (max = 90deg/s) speeds in a virtual environment. This virtual world comprised a ground plane whose textural elements had limited lifetime (250 ms) to avoid serving as landmarks. The ground plane was circular with a radius of 70 m (near and far clipping planes at 5 cm and 40 m respectively), with the subject positioned at its center at the beginning of each trial. Each texture element was an isosceles triangle (base x height: 8.5 x 18.5 cm^2^) that was randomly repositioned and reoriented at the end of its lifetime, making it impossible to use as a landmark. The standard density of the ground plane elements (i.e., the density on which animals were trained) was 5.0 elements/m^2^ (high-density condition). This density was reduced by a factor of 50 (0.1 elements/m^2^) in a manipulation condition to force the animals to further use their acquired internal model, rather than solely sensory evidence. All stimuli were generated and rendered using C++ Open Graphics Library (OpenGL) by continuously repositioning the camera based on joystick inputs to update the visual scene at 60 Hz. The camera was positioned at a height of 0.1 m above the ground plane. Spike2 software (Cambridge Electronic Design Ltd., Cambridge, UK) was used to record and store the timeseries of target and animal’s location, animal linear and angular velocity, as well as eye positions. All behavioral data were recorded along with the neural event markers at a sampling rate of 833.33Hz.

### Behavioral Task

Monkeys steered to a target location (circular disc of radius 20 cm) that was cued briefly (300ms) at the beginning of each trial. Each trial was programmed to start after a variable random delay (truncated exponential distribution, range: 0.2 – 2.0 s; mean: 0.5 s) following the end of the previous trial. The targets appeared at a random location between –45 to 45 deg of visual angle, and between 1 and 4 m relative to where the subject was stationed at the beginning of the trial. The joystick was always active, and thus monkeys were free to start moving before the target vanished, or before it appeared. Monkeys typically performed blocks of 750 trials before being given a short break. In a session, monkeys would perform 2 or 3 blocks. High (5.0 elements/m^2^) and low (0.1 elements/m^2^) density conditions were randomly intermixed with equal probability within each block. Feedback, in the form of juice reward was given following a variable waiting period after stopping (truncated exponential distribution, range: 0.1 – 0.6 s; mean: 0.25 s). They received a drop of juice if their stopping position was within 0.6 m away from the center of the target (“reward boundary”). No juice was provided otherwise. Monkeys were trained by gradually reducing the size of the reward zone until their performance stopped improving.

### Neural Recording and Pre-Processing

We recorded extracellularly, either acutely using a 24-channel linear electrode array (100 μm spacing between electrodes; U-Probe, Plexon Inc, Dallas, TX, USA; MSTd in Monkey Q and S, and dlPFC in monkey M) or chronically with multi-electrode arrays (Blackrock Microsystems, Salt Lake City, UT, USA; 96 electrodes in area 7a in Monkey Q, and 48 electrodes in both area 7a and dlPFC in monkey S; see Figure 2 – supplement 1 in Ref^35^ for pictures and reconstruction of these recording devices in brain). Chronic setups were used for their higher yield, while acute linear probe recordings were used in order to target MSTd, which is deep. During acute recordings with the linear arrays, the probes were advanced into the cortex through a guide-tube using a hydraulic microdrive. Spike detection thresholds were manually adjusted separately for each channel to facilitate real-time monitoring of action potential waveforms. Recordings began once waveforms were stable. The broadband signals were amplified and digitized at 20 KHz using a multichannel data acquisition system (Plexon Inc, Dallas, TX, USA) and were stored along with the action potential waveforms for offline analysis. Additionally, for each channel, we also stored low-pass filtered (-3dB at 250Hz) local-field potential (LFP) signals. For the array recordings, broadband neural signals were amplified and digitized at 30 KHz using a digital headstage (Cereplex E, Blackrock Microsystems, Salt Lake City, UT, USA), processed using the data acquisition system (Cereplex Direct, Blackrock Microsystems) and stored for offline analysis. Additionally, for each channel, we also stored low-pass filtered (-6 dB at 250 Hz) local-field potential (LFP) signals sampled at 500 Hz. Finally, copies of event markers were received online from the stimulus acquisition software (Spike2) and saved alongside the neural data.

Spike detection and sorting were initially performed on the raw (broadband) neural signals using KiloSort 2.0 software on an NVIDIA Quadro P5000 GPU. The software uses a template matching algorithm both for detection and clustering of spike waveforms. The spike clusters produced by KiloSort were visualized in Phy2 and manually refined by a human observer, by either accepting or rejecting KiloSort’s label for each unit. In addition, we computed three isolation quality metrics; inter-spike interval violations (ISIv), waveform contamination rate (cR), and presence rate (PR). ISIv is the fraction of spikes that occur within 1ms of the previous spike. cR is the proportion of spikes inside a high-dimensional cluster boundary (by waveform) that are not from the cluster (false positive rate) when setting the cluster boundary at a Mahalanobis distance such that there are equal false positives and false negatives. PR is 1 minus the fraction of 1 minute bins in which there is no spike. We set the following thresholds in qualifying a unit as a single-unit: ISIv < 20%, cR < 0.02, and PR > 90%.

### Analyses

#### Behavior

The location of targets and the monkey’s end locations were expressed in polar coordinates, with a radial distance (target = *r*, response = 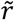) and eccentricity from straight ahead (target = *θ*; response = 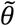). On a subset of trials (∼13%) animals stopped after <0.5m, suggesting they aborted the trial. Similarly, on a subset of trials (∼5%) animals did not stop during the course of a trial (max duration = 7 seconds). These trials were discarded before further behavioral analyses. A linear model with multiplicative gain accounted well for the observed data (average R^2^ = 0.72). Thus, we used the slopes of the corresponding linear regressions as a measure of bias (**Fig. 1B** and **C**). We also report the fraction of trials in which the animal was rewarded (reward radius = 65cm centered on the firefly, determined by staircase such that animals would be rewarded on ∼66% of trials). Lastly, to index variance in endpoint responses we compute for each session the S.E.M of distances to target (rendering a two-dimension error, along x and y, into a single dimension, distance). We then report the mean S.E.M and variance across sessions and density conditions.

#### Generalized Additive Model

The Poisson Generalized Additive Model (P-GAM) defines a non-linear mapping between spike counts of a unit *y*_*t*_ ∈ ℕ_0_ (binned at 6ms temporal resolution) and a set of continuous covariates ***x***_*t*_ (angular and linear velocity and acceleration, angular and linear distance traveled, angular and linear distance to target, and LFP instantaneous phase across different frequency ranges), and discrete events ***z***_*t*_ (time of movement onset/offset, target onset, reward delivery, and the spike counts from simultaneously recorded units). A unit’s log-firing rate was modeled as a linear combination of arbitrary non-linear functions of the covariates,

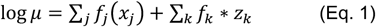

where * is the convolution operator, and the unit spike counts are generated as Poisson random variables with rate specified by (Eq. 1). Input specific non-linearities *f*(⋅) are expressed in terms of flexible B-splines, *f*(⋅) ≈ ***β*** ⋅ ***b***(⋅) and are associated with a smoothness enforcing penalization term controlled by a scale parameter *λ*_*f*_,

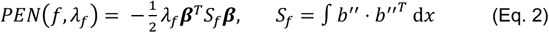

The larger *λ*_*f*_, the smoother the model. This penalization terms can be interpreted as (improper) Gaussian priors over model parameters, the resulting log-likelihood of the model takes the form,

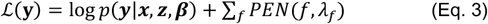

with ***y*** ∈ ℝ^***T***^ the spike counts of the unit, ***x*** ∈ ℝ^*J*×*T*^ and ***z*** ∈ ℝ^*K* ×*T*^, *T* the time points, ***β*** the collection of all B-spline coefficients and *p*(⋅) the Poisson likelihood. Both parameters ***β*** and the hyperparameters ***λ*** are learned from the data by an iterative optimization procedure that switches between maximizing Eq. 3 as a function of the parameters, and minimizing a cross-validation score as a function of the hyperparameters (see^30^ for further details). The probabilistic interpretation of the penalization terms innovatively allowed us to compute a posterior distribution for the model parameters, derive confidence intervals with desirable frequentist coverage properties, and implement a statistical test for input variable inclusion that selects a minimal subset of variables explaining most of the variance.

#### Coupling Filters

Coupling filters (and the corresponding inferential statistics) were determined via the P-GAM^30^. Within area coupling filters were set to a duration of 36ms, and across area filters were initially set to a duration of 600ms. We focus our analyses exclusively on the within area coupling filters, given their higher presence rate, and the fact that we can be more confident in indexing unit-to-unit interactions when restricting the analyses to a short time-frame (i.e., 36ms). For the coupling probabilities reported in **Fig. S2D**, we corrected for the effects of unit distance (i.e., **Fig. S1A**) by first fitting a brain region-specific logistic regression. Specifically, we expressed coupling probability as a non-linear function of electrode distance as follows,

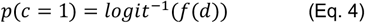

With *c* being a binary variable taking value 1 for significant coupling and 0 otherwise, *d* being the electrode distance, and *f* being a non-linear function expressed in terms of B-splines. Each brain area was fit independently, and the coupling probability in **Fig. S2D** was set as the model prediction for a distance of 500um. Coupling probability was computed as the fraction of significantly coupled units within area pairs (p-value <0.001 covariate inclusion test). Coupling strength (i.e., **Fig. 2C**) was computed as the norm of the betas defining the coupling filter. Coupling stability (e.g., **Fig. 2D**) was computed as the correlation between coupling filters in the low- and high-density condition. Chance and ceiling levels were established by permutation. To relate the change in within area functional connectivity with the change in behavior caused by the decrease in optic flow signals, we computed the average coupling stability per area and session, given the session had at least 5 coupled units and a P-GAM pseudo-R^2^ greater than 0.01.

#### Latent Population Dynamics

We computed population latent dynamics by principal component analysis (PCA). For the illustration in **Fig. 3A**, PCA was based on trial averaged firing rates in rewarded trials across all sessions (i.e., ‘pseudo-populations). We segmented trials between the time of target onset and movement offset, and estimated instantaneous firing rates by convolving spike counts (binned at 6ms) with a 100ms Gaussian window. We matched trial durations by linearly interpolating the rates (i.e., time-warping). Then, for the visualization, we grouped trials into eighteen categories (9 target positions x 2 densities; see **Fig. 3A**, top) and averaged the instantaneous rate within each category. We then stacked all rates (for all time points and categories) in a single vector ***r***_*i*_ ∈ ℝ^*T*⋅*C*⋅*D*^, where *i* = 1, …, *N* is the neuron index, T is the number of the interpolating time points, C is the target location (categorized), and D are the density conditions. We computed the PCA projection based on the (T ⋅ C ⋅ D) × N matrix of the population rates, and finally projected each condition separately on the PCA axis. To quantitatively assess the stability of global population dynamics across density conditions we developed a permutation test that compares the dynamics of similar trials within and between density conditions. For each session we first extracted the PCA axis by stacking the instantaneous firing rate of all rewarded trials (estimated as above). Subsequently, we applied the following algorithm 1000 times per session: 1) randomly select a reference trial from the high-density condition; 2) selected two trials for comparison, a high-density trial and a low-density trial, for which the initial target location was less then 10cm away from that of the reference trial; 3) project the population activity of these 3 trials onto the PCA axis; 4) compute the Frobenius norm-based distance between the projected activity of the target trial and each of the two comparison trials. This procedure yielded the distributions shown in **Fig. S4A** (5 example sessions shown). Finally, we computed the d-prime score separating these distributions (**Fig. S4B**). This procedure was repeated while projecting neural activity on 1 to 10 PCs. Similarly, we repeated this analysis while conducting PCA separately on the different density conditions, again yielding conceptually identical results.

#### Decoding, Lasso Regression

To specifically examine the manifold coding for distance to target, we decoded this variable (the critical task-relevant latent variable, given that the target is invisible) from the population of instantaneous firing rates estimated by convolving spike counts with a 100ms Gaussian window. For a fair comparison of the decoding accuracy between areas, we matched the number of units and the mean firing rate for each area on a session-by-session basis. Then, we split trials according to the density condition and, within density, we used 90% of trials for training a linear decoder. The remaining 10% of data was used to estimate the decoder prediction accuracy. For cross-context decoding we trained on high-density optic flow and decoded on the low density condition. We opted for a Lasso regularized regression to control for overfitting. Lasso hyperparameters were selected by grid-search with a 5-fold cross-validation scheme. Model prediction accuracy within and across density condition was quantified as cross-validated r^2^ (**Fig. 3B**, example distances to target and their reconstruction shown in **Fig. 3C**). To test for the role of neural co-fluctuations in allowing for distance to target decoding, we adopted a shuffling procedure that destroyed neuron to neuron correlations but kept signal correlations intact. Namely, we first discretized distance to target in 15 bins (other discretization tested and yielded conceptually identical results). Then, since each time-point corresponded to a particular bin, for each neuron we shuffled the counts for each time point corresponding to the same bin. This procedure leaves tuning functions unchanged (as we are shuffling like-for-like) but disrupts the correlation between neurons. We then conducted the lasso regression as detailed above. This shuffling procedure was done 10 times per neuron and session to assure reproducibility (**Fig. 3D** plots the first run, after checking results were reproduced across all shuffling runs).

## Acknowledgements

The authors thank Jing Lin and Jian Chen for programming the experimental stimulus. We also thank Eric Avila, Kaushik Lakshminarasimhan, Stefania Bruni, and Panos Alefantis for data collection, and Roozbeh Kiani for his surgical expertise during the Utah array implantations. The work was funded by 1U19 NS118246, 1R01 NS120407, and 1R01DC014678 from NIH to D.E.A, by 1R01MH125571 from NIH, the National Science Foundation under NSF Award No. 1922658, and a Google faculty award to C.S, and by K99NS128075 to JPN.

## Supplementary Figures and Figure Captions

**Figure S1.**
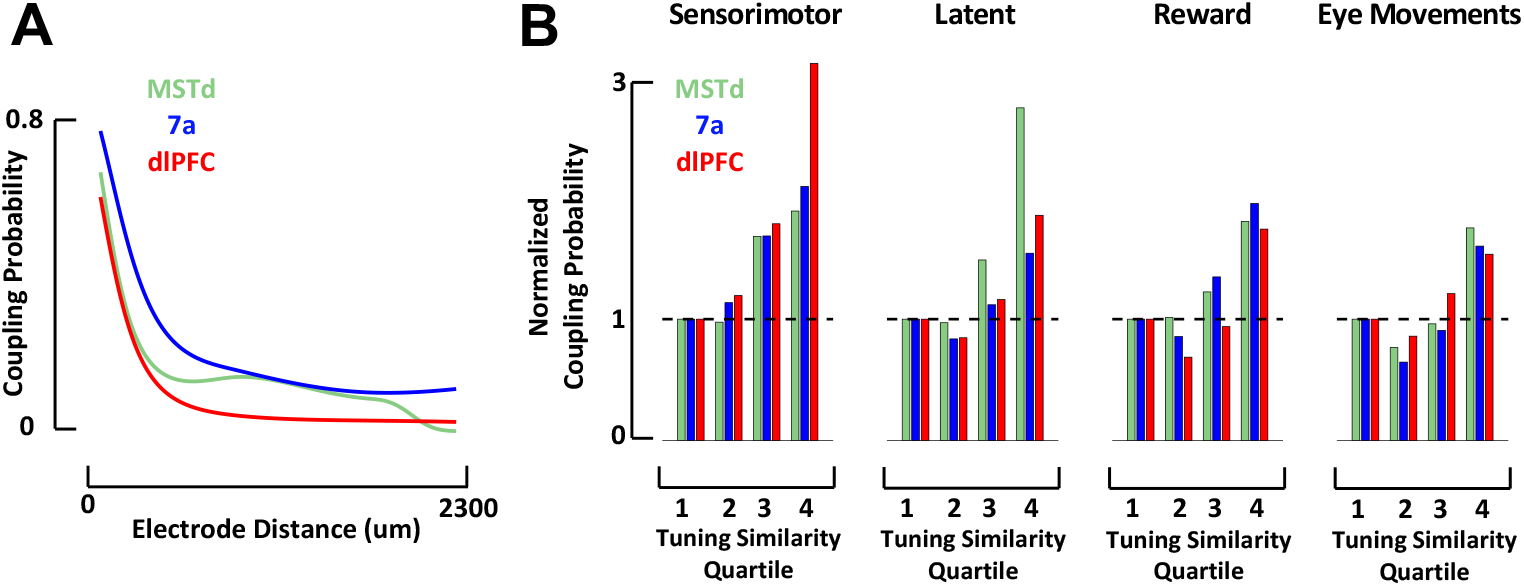
**A**. Coupling probability (y-axis) between two neurons as a function of the distance between them (x-axis). This illustration is for data from a single monkey (Monkey S) with recordings in MSTd (green, number of coupling pairs = 1714), area 7a (blue, number of coupling pairs = 97711), and dlPFC (red, number of coupling pairs = 41665). As expected, neurons that are closer to each other are more likely to be coupled. Given this effect and that multiple recording techniques were used (with different spacing between electrodes, 400um in Utah array and 100um in linear probes), we used these estimates to correct coupling probabilities to a single distance (500um). **B**. Coupling probability between any two units given their tuning similarity to sensorimotor, latent, or other (reward and eye positions) variables (data from 3 monkeys). Tuning similarity is computed as the correlation in tuning functions, then these are discretized in quartiles by their tuning similarity (r^2^ = [0-.25; 0.25-0.5; 0.5-0.75; 0.75-1]) and averaged within each category (e.g., sensorimotor or latent) and tuning similarity bin. The coupling probability is expressed as a ratio, normalized to the bin with lowest tuning similarity (leftmost), such that a normalized coupling probability of ∼3 (e.g., sensorimotor variables in dlPFC) indicates that coupling is three times more likely given high vs. low tuning similarity.

**Figure S2.**
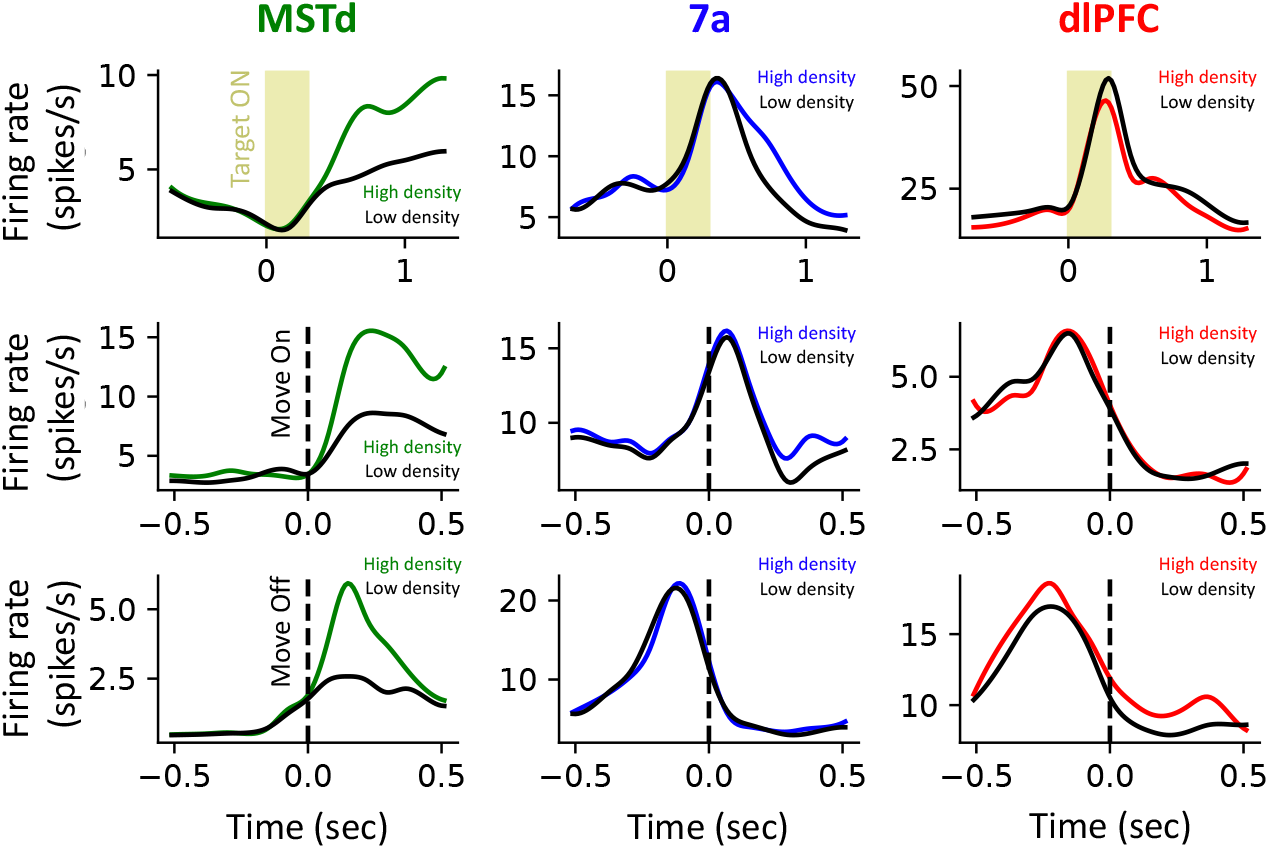
Example peri-event time histograms (PETH). Nine example PETH, three per brain area (MSTd, 7a, and dlPFC, respectively in green, blue, and red) and per event; target onset (top row), macaque movement onset (middle row) and macaque movement offset (bottom row). Most importantly, the PETHs are plotted separately for the high density optic flow condition (colored by brain area) and the low density optic flow conditions (black). The examples are representative (see main text and **Figure 2B**) in that MSTd is gain modulated (gain <1) from the high to low density condition, while responses in 7a and dlPFC were less impacted by the density of optic flow.

**Figure S3.**
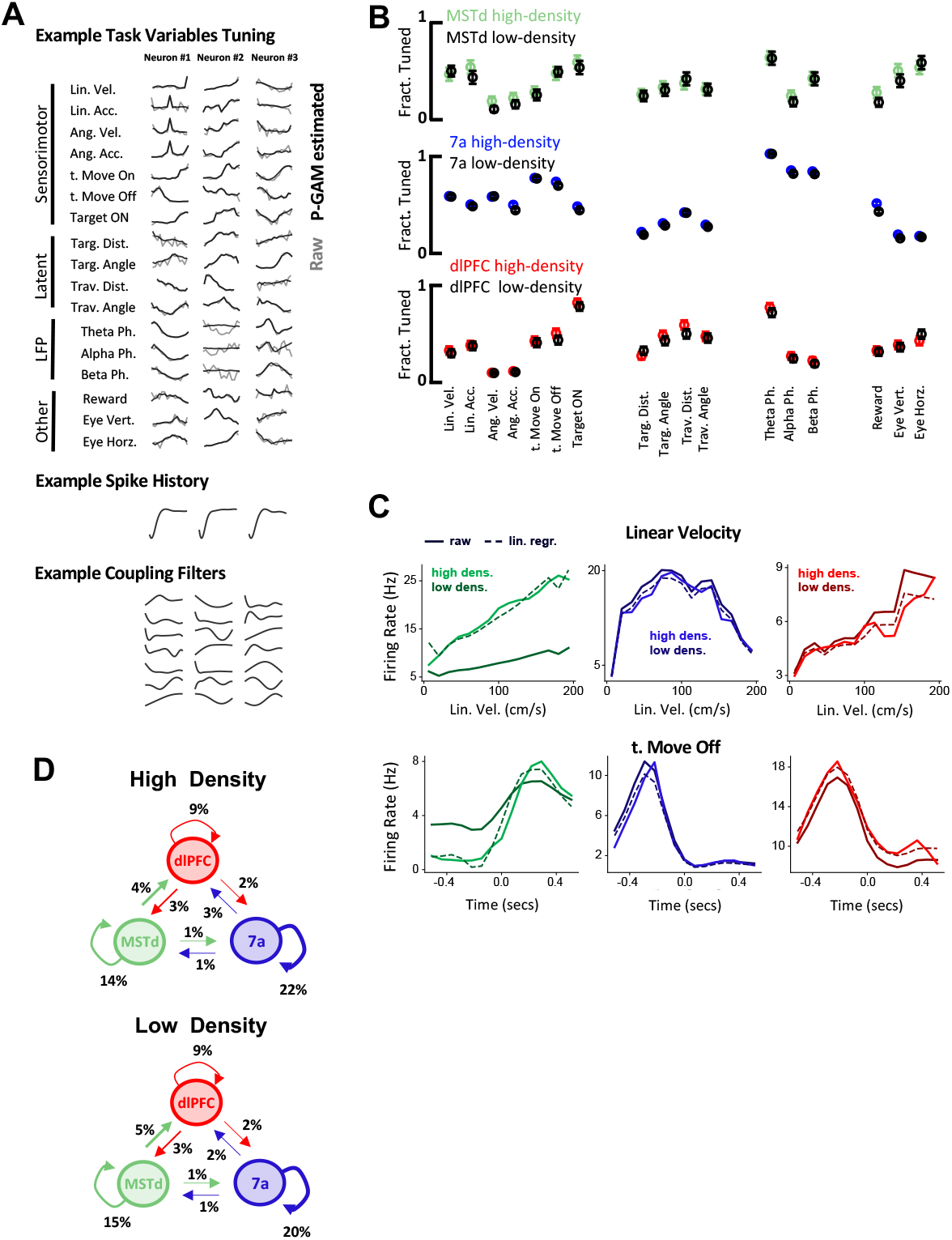
**A**. Example filters estimated by the P-GAM. From top to bottom: Raw (gray) and P-GAM reconstructed (black) filters for task-variables (demonstrating the model’s ability to account for spiking activity); Spike history filters show the characteristic refractory period of single units; Coupling filters (see Figure S3 for more examples). **B**. Fraction of neurons tuned. High-(colored) and low-density (black) optic flow conditions did not result in a different fraction of neurons tuned to different task variables in MSTd (green), area 7a (blue), or dlPFC (red). Error bars are +/-95%CI across neurons in all sessions. **C**. Example tuning functions to linear velocity (top) and time from movement offset (bottom) for each neural area (green = MSTd; blue = area 7a; red = dlPFC) during high-(brighter color) and low-(darker color)) optic flow density conditions. In addition to the raw data (solid lines), we also demonstrate the re-scaling from low- and high-density condition (dashed lines) in order to estimate gain modulations. The examples are representative, in that they demonstrate no gain modulation in 7a and dlPFC, but a strong modulation in MSTd. **D**. Fraction of units coupled within and across areas in high- and low-density conditions. Thickness of arrows, and inset percentage indicate the fraction of neurons coupled within and across areas. An arrow projecting from e.g., MSTd to dlPFC indicates that the firing of a neuron in MSTd will subsequently influence spiking activity in dlPFC.

**Figure S4.**
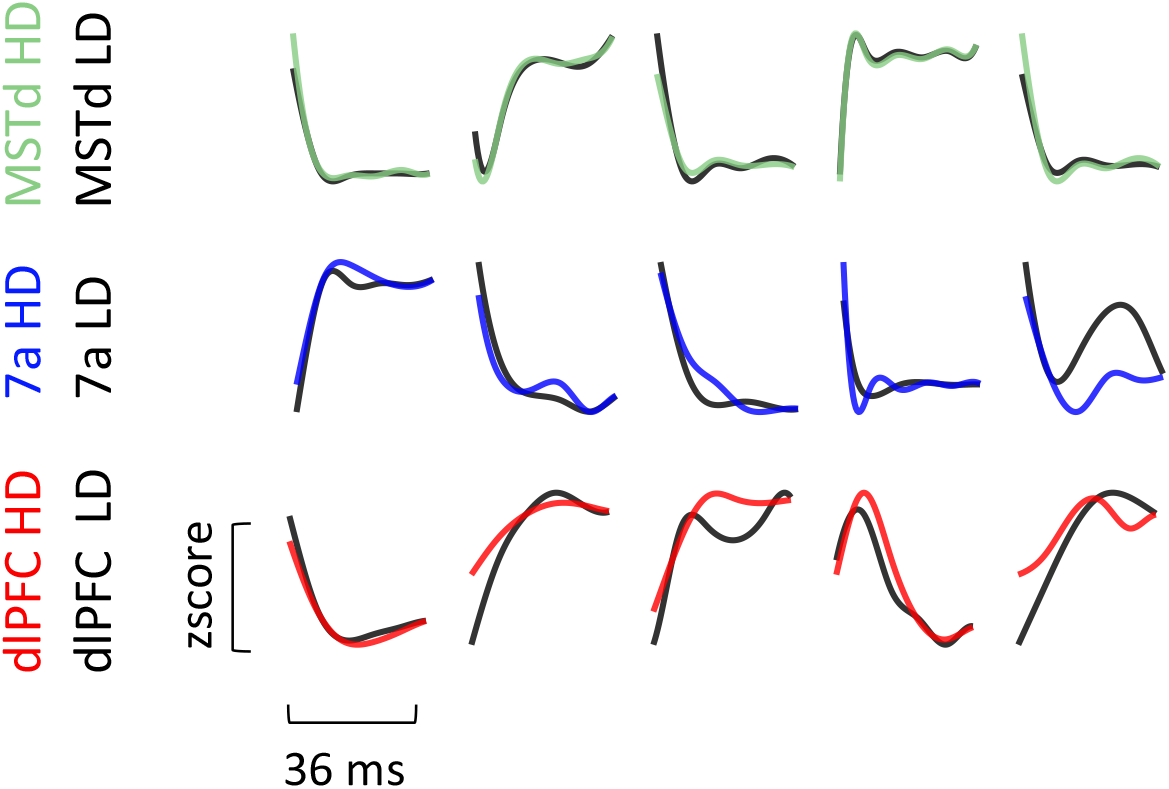
Example coupling filters. Coupling filters remain stable across optic flow densities in MSTd (green), but less so in area 7a (blue) and dlPFC (red). Five examples are shown for each area (high-density colored and low-density in black). Coupling functions within area had a length of 36ms (see^16^ and Methods for further detail).

**Figure S5.**
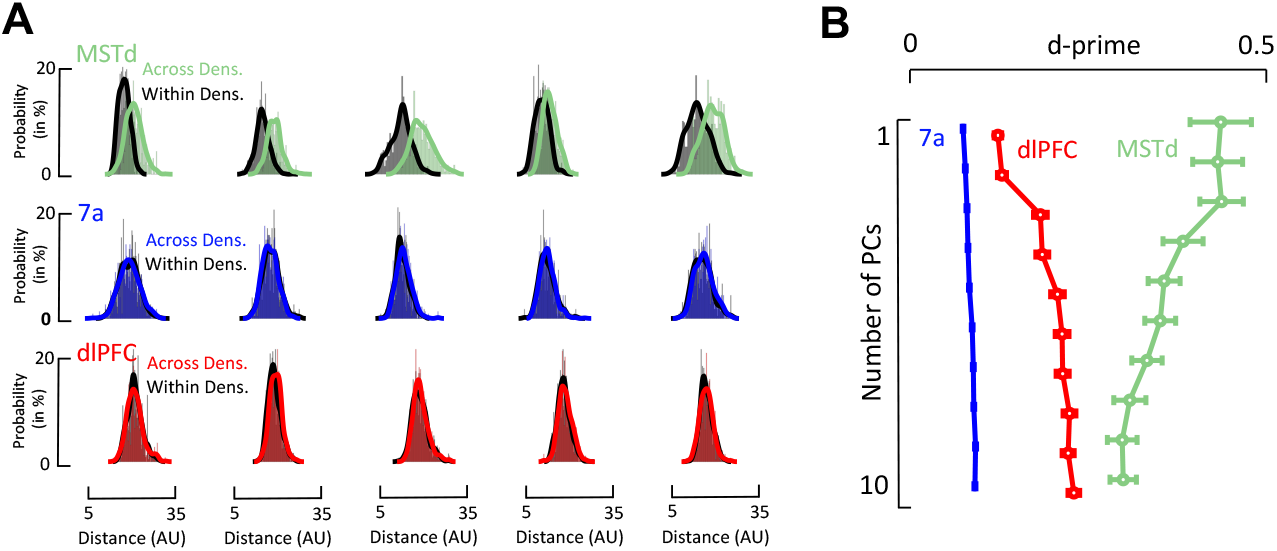
Population codes remain stable in 7a and dlPFC but not MSTd. A. Distribution of distances between high- and low-density conditions in latent space. Five example sessions are shown for MSTd (green), 7a (blue) and dlPFC (red). In each session, 1000 trials are paired based on their similarity in target location and steering behavior, either within (i.e., HD/HD or LD/LD) or across (i.e., HD/LD) density conditions. Each trial is then projected onto PC space (2D) and their distance is computed. The histograms show are the distances in latent space between those 1000 trials. **B. Summary statistics**. The distance between distributions (within vs. across, as in **A**) is computed by d-prime. Latent neural trajectories are more dissimilar across density conditions in MSTd (green) than 7a (blue) or dlPFC (red) regardless of the number of PCs used. As we index higher-order PCs, we observe that dlPFC become different between high- and low-density conditions. Area 7a is always stable. Error bars are ± 1 S.E.M.

